# Field margins as substitute habitat for the conservation of birds in agricultural wetlands

**DOI:** 10.1101/2022.05.05.490780

**Authors:** Mallet Pierre, Béchet Arnaud, Sirami Clélia, Mesléard François, Blanchon Thomas, Calatayud François, Dagonet Thomas, Gaget Elie, Leray Carole, Galewski Thomas

## Abstract

Breeding birds in agricultural landscapes have declined considerably since the 1950s and the beginning of agricultural intensification in Europe. Given the increasing pressure on agricultural land, it is necessary to identify conservation measures that consume little productive land. We tested the compensation hypothesis which states that field margins may represent substitute habitats for bird species in agricultural wetlands. We monitored bird species in 86 crop fields in rice paddy landscapes of Camargue (southern France), a wetland of international importance for birds. We investigated whether the area of three types of field margins (reed strips, grass strips and hedgerows) within a 500 meter buffer around the centroid of each crop fields had an effect on the abundance of bird species from three groups defined based on their primary habitat (reedbeds, grasslands, and forest edge species). We controlled for the area of each type of semi-natural habitat (wetlands, grasslands, and woodlands), crop diversity (rice, wheat, alfalfa, rape, and market gardening) and mean crop field size. Results show partial support of the compensation hypothesis with species-dependent responses to primary and substitute habitat area. Some species within the reedbed and grassland bird guilds are favored by the area of their primary habitat as well as by the area of field margins, in line with the compensation hypothesis. Eurasian reed warbler is favored by the area of both wetlands and reed strips. Corn bunting is favored by grassland and grass strip areas. We could not confirm the compensation hypothesis for other species. However, this may be due to the fact that most of these species did not respond to their primary habitat. These results therefore suggest that field margins may represent substitute habitats for some species but further studies, in contexts where species are strongly associated with their primary habitat, would be needed to confirm the generality of this hypothesis. Our results also suggest that species response to increasing the area of a field margin type may vary among guilds and even within guilds. Therefore, it may be difficult to favor all species within a given landscape and management actions may need to be tailored to whichever species are locally associated with the highest conservation priority. To tackle this challenge, it may be necessary to design landscape management actions at different spatial scales.

## 1 Introduction

Farmland bird populations have experienced a massive decline worldwide in recent decades, primarily due to the loss of semi-natural habitats and intensification of agricultural practices (PECBMS, 2022; Stanton et al., 2018; Sundar and Subramanya, 2010). Agricultural areas represent 37 % of the European terrestrial area and host a large proportion of terrestrial biodiversity (DataBank, 2018; Herzog et al., 2013). It is therefore not practical to rely solely on the creation of protected areas to compensate for the declines in biodiversity observed in European agricultural environments (Meyer et al., 2013; Warren et al., 2021). Rather, conservation efforts should also focus on maintaining and increasing the capacity of agricultural landscapes to support biodiversity through the adoption of biodiversity-friendly agricultural practices and the protection of non-productive refuge areas, i.e. promote land sharing (Grass et al., 2021).

Patches of semi-natural habitats, such as woodlands, grasslands and wetlands, remaining within agricultural landscapes may provide permanent habitat for wildlife and host a large part of farmland biodiversity (Newton, 2017; Toffoli and Rughetti, 2017). However, these patches are scarce and under increasing pressure in Europe due to agricultural intensification which leads to their progressive conversion to arable land despite efforts from the European Union to slow down this trend through agri-environment schemes (Batáry et al., 2015). Hence, in some agricultural landscapes, field margins, i.e. linear elements covered by semi-natural vegetation along the edge of crops, are the only type of semi-natural habitat left (Marshall and Moonen, 2002). The habitat compensation hypothesis states that species may compensate for the loss of their primary habitat by using alternative habitats as a substitute (Norton et al., 2000). For instance, Montagu’s harrier (*Circus pygargus*) primarily nests in shrublands and grasslands but, in some part of its distribution range, it now relies exclusively on crop fields for breeding and foraging (Norton et al., 2000). It has also been shown that aquatic invertebrates can use drainage ditches as substitute habitats for natural lakes and rivers (Dollinger et al., 2015). The habitat compensation hypothesis has been investigated in the context of farmland abandonment and in dry agricultural areas (e.g. Brotons et al., 2005; Saura et al., 2014; Vallecillo et al., 2008) but rarely in wetland agricultural areas (e.g. Decleer et al., 2015) despite their specific landscape characteristics and biodiversity.

One of the main crops cultivated in wetlands is rice, a flooded cereal which represents 22.8 % of the world cereal surface area (FAO, 2018; Singh et al., 2001). In such rice paddy landscapes, agricultural and semi-natural areas are generally intermingled with the presence of large field margins along ditches. Among birds associated with these rice paddy landscapes, there are both waterbirds (e.g. gulls, terns, herons, storks, ibises, waders…) and terrestrial bird species. While the role of rice paddy landscapes as alternative habitat for waterbirds has been largely studied, their role for terrestrial birds has received much less attention (Elphick, 2015). Considering the long-term decline of terrestrial bird populations in agricultural landscapes (Fraixedas et al., 2019), identifying conditions favoring them would be useful to improve recommendations for agri-environmental management practices in rice paddy landscapes. Terrestrial birds using rice paddy landscapes include different ecological guilds: reedbed birds, which are primarily associated with freshwater marshes (Morganti et al., 2019); forest edge species, which are originally associated with forest borders and clearings (Hinsley and Bellamy, 2019; Newton, 2017); and grassland species, which originally live in grassy or shrubby vegetation with no tree cover (Di Giacomo et al., 2010). Field margins could provide resources and nesting habitats for these species (Vickery et al., 2009), e.g. reed strips along ditches for reedbed birds, hedgerows for forest edge species and grass strips for grassland species. However, the role of field margins for terrestrial birds has rarely been considered in studies conducted in rice paddy landscapes (King et al., 2010).

The Camargue (Rhône delta) is a biologically rich area listed in the Ramsar Convention and classified as a Biosphere Reserve by UNESCO (Blondel et al., 2019). Natural areas cover 58,000 ha and agricultural areas 55,100 ha (Tamisier and Grillas, 1994). Rice represents 48 % of the crop area and is mainly cultivated in rotation with wheat (19 %) and alfalfa (5 %). Within this region, bird species associated with agricultural areas have experienced the greatest rate of decline over the past 50 years compared to waterbirds (Fraixedas et al., 2019; Galewski and Devictor, 2016). Hence, it is critical to assess whether field margins could constitute a lever for bird conservation as their restoration and management may be readily changed by farmers.

In this paper, we tested the habitat compensation hypothesis in rice paddy landscapes of Camargue by assessing whether field margins act as substitute habitats for reedbed birds, forest edge birds and grassland birds. We conducted bird surveys in 86 crop fields in Camargue. Specifically, we predicted that (i) forest edge species would be positively influenced by woodlands and hedgerows; (ii) grassland birds would be positively influenced by grasslands and grass strips and (iii) reedbed birds would be positively influenced by semi-natural wetland areas and reed strips.

## 2 Material and methods

### 2.1 Study area

Our study was conducted in the Rhône River delta, a 180,000 ha polderized flood plain located in Southern France and known as “Camargue”. Warm summers typical of the Mediterranean climate (average monthly temperature between May and October above 15°C; Blondel et al., 2019), as well as fresh water pumped from the Rhône River allows rice cultivation. This flooded crop is essential for washing out salt-rich soils and allows rotation with dry crops, mainly wheat and alfalfa. In Camargue, field margins are often wide (> 3 m) to be waterproof and keep the crop fields flooded during the rice cultivation period. Several types of vegetation can therefore co-occur within the same field margin, such as reed strips, hedgerows or grass strips. In Camargue, the area of semi-natural habitats decreased from 67 % to 39 % between 1942 and 1984 and since remained stable at around 58,000 ha (Mallet, 2022; Tamisier and Grillas, 1994). These semi-natural areas are spatially segregated in the delta; woodlands are mainly restricted to riparian areas along the Rhône River, wetlands occupy depressions and cover large areas in the center and south of the delta while grasslands (mostly constituted of meadows and salt steppes) surround the wetlands on slightly elevated areas (Appendix A).

### Study design

We selected 86 crop fields belonging to 17 farms across the Camargue (Fig. 1). All fields were organic to limit confounding effects associated with variation in the intensity of agricultural practices. We selected crop fields covered by the crop types representative of the main agricultural production in Camargue (rice, wheat, alfalfa, rape, and market gardening). Crop fields were selected along two independent gradients of semi-natural cover and hedgerow cover using the methodology developed by Pasher et al., (2013). To do so, we measured semi-natural and hedgerow areas in a 500 meters square moving window with a step size of 100 meters around every agricultural land of Camargue thanks to land-use data from 2019 of the BD TOPO^®^, OSO Land Cover Map and the Regional Natural Park of the Camargue. No maps of grass strip or reed strip were available prior to crop field selection. Therefore, we checked for the distribution of sampled crop fields along gradients of explanatory variables once the selection and on-site mapping were completed. We also checked for correlation among the cover of different types of field margin and other landscape variables (see below).

**Figure 1.**
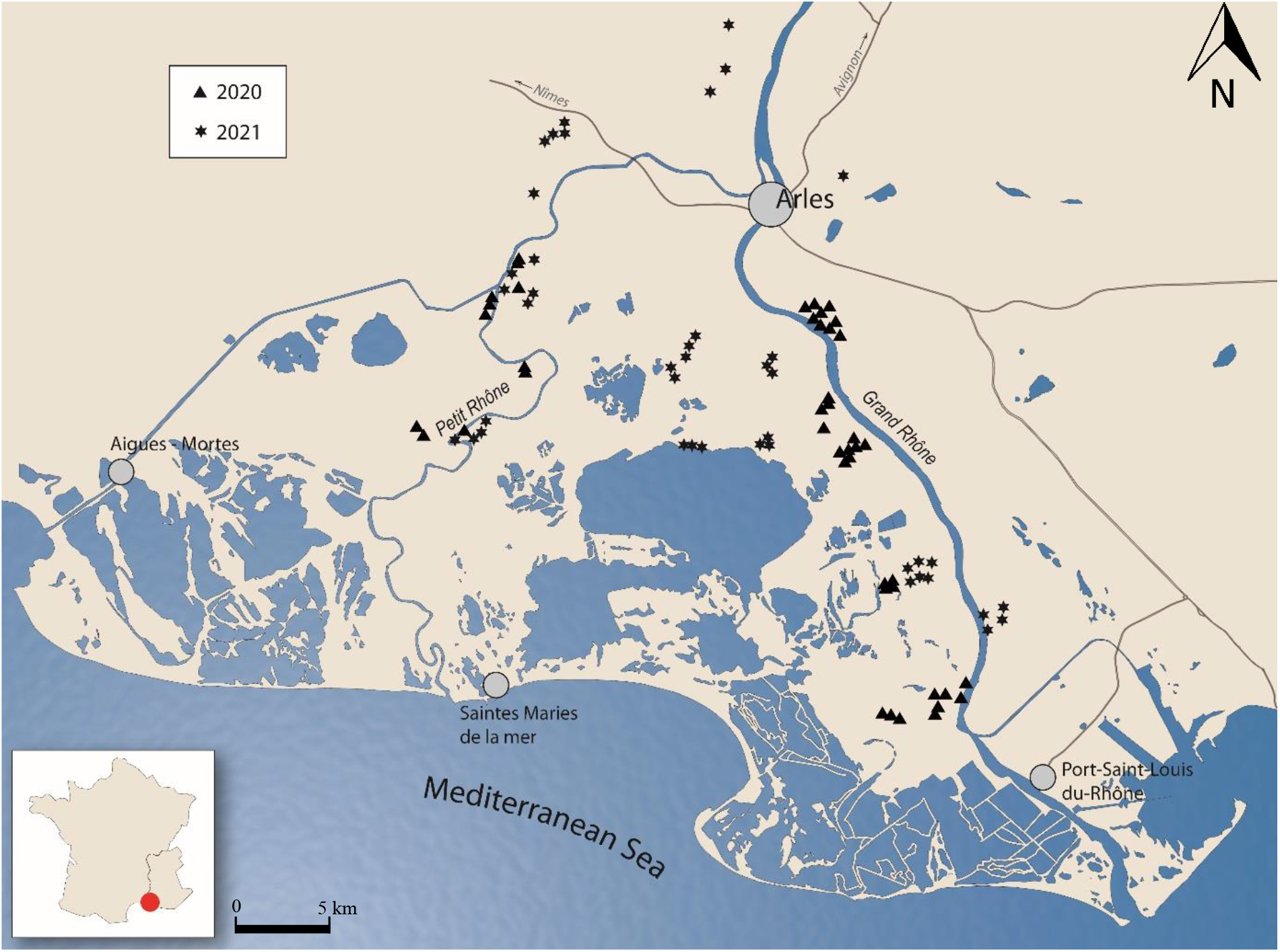
Location of the 86 crop fields monitored for birds in Camargue, Rhône delta. Triangles represent crop fields sampled in 2020 and stars represent crop fields sampled in 2021.

We calculated the area of each type of field margin and semi-natural habitat within a 500 meter buffer around the centroid of each crop field, following Chan et al. (2007). The maximum size of the sampled crop fields was 14 ha, hence much smaller than this buffer. First, we estimated the area of the three types of field margins: (1) hedgerows, tree lines and bushy areas; (2) grass strips, grassy boundaries including grassy tracks or dirt roads used for the moving of agricultural machinery; (3) reed strips that grow in and along irrigation or drainage earthen ditches. Because we aimed at testing the hypothesis that field margins represent substitute habitats whatever their shape, we calculated the area and not the length of field margins. Second, we estimated the area of three categories of semi-natural areas: (1) woodlands (mainly riparian forests dominated by white poplar (*Populus alba*), pinewoods (*Pinus pinaster*), and tamarisk (*Tamarix gallica*) groves) and shrublands dominated by narrow-leaved mock privet (*Phillyrea angustifolia*); (2) grasslands including dry grasslands extensively grazed by free-range cattle, Mediterranean salt meadows and halophilous scrubs and fallow lands; (3) wetlands including freshwater and brackish marshes, reedbeds and ponds. Landscape mapping was based on field observations done after the bird monitoring in June 2020 and June 2021 (see below) because fine scale assessment was not feasible based on remote sensing approaches only, particularly for reed strips. Finally, to account for the possible confounding effect of crop field heterogeneity, we also estimated within each 500 meter buffer the mean crop field size and the Shannon diversity index of crop types (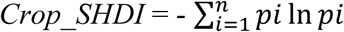, where *pi* corresponds to the proportion of crop cover type *i* in the landscape), following the method implemented in Sirami et al. (2019). As a result, we obtained values for eight landscape variables for each sampled crop field.

### 2.3 Bird monitoring and traits

Birds were monitored over 5-minute point counts halfway along the longest field margin of each crop field during the breeding period. Two visits were conducted at each site between mid-April and mid-June with at least 4 weeks between the two visits, following the protocol from the French common breeding bird census scheme (Jiguet, 2003). Flying birds were removed from the analyses because they were not interacting directly with the landscape. Birds landing outside the sampled crop field and its field margins were also removed to avoid detection bias potentially generated by hedgerows preventing the observer to see birds beyond trees. We used the maximum abundance per site between the two visits for each species for further analyses.

We assigned each species to one of three guilds according to the primary habitat used for breeding: reedbed, grassland and forest edge birds. Assignment was based on the EUNIS habitat classification that describes species communities related to woodlands, wetlands, grasslands or urban areas (Appendix B). Generalist birds, i.e. not linked to one habitat in particular, or birds that use urban areas for breeding were discarded from the analyses. We modulated the EUNIS data with information provided by a local expert (T.G.) to take into account ecological particularities of the Camargue. To avoid extreme cases of zero-inflation, we only kept species present in more than 15 % of the sampled crop fields (Marja and Herzon, 2012). Fourteen species were retained for the analyses (Table 1).

**Table 1.** Species studied within the three guilds based on the EUNIS database combined with information provided by local experts to take into account ecological particularities of the Camargue (Appendix B).

| Guilds | Species |
| --- | --- |
| Forest edge birds | European greenfinch (*Chloris chloris*) |
|  | Carrion crow (*Corvus corone*) |
|  | Melodious warbler (*Hippolais polyglotta*) |
|  | Common nightingale (*Luscinia megarhynchos*) |
|  | Great tit (*Parus major*) |
|  | Eurasian magpie (*Pica pica*) |
|  | European green woodpecker (*Picus viridis*) |
|  | Eurasian blackcap (*Sylvia atricapilla*) |
|  | Eurasian hoopoe (*Upupa epops*) |
| Grassland birds | Crested lark (*Galerida cristata*) |
|  | Corn bunting (*Emberiza calandra*) |
|  | Eurasian skylark (*Alauda arvensis*) |
| Reedbed birds | Eurasian reed warbler (*Acrocephalus scirpaceus*) |
|  | Great reed warbler (*Acrocephalus arundinaceus*) |

In order to check for the completeness of our data, we calculated the coverage of our sampling, which is defined as the proportion of the total number of individuals in an assemblage that belong to the species present in the sample (Chao and Jost, 2012). This index corresponds to the probability of occurrence of the species observed in the sample. The coverage was calculated by crop field for all 14 species considered within the present study. The overall coverage of our sampling was 73.5 %, which reflects no undersampling issue (Mallet et al., 2022). The sampling completeness per crop field was not correlated with any explanatory variable (Pearson coefficient < 0.24, Appendix C), which suggests that the study design was robust and not biased toward one or several landscape variables.

### 2.4 Data analysis

We ran one linear mixed-effect model with bird abundance as the response variable, while fixed effects were species identity, the area of the three field margin types (hedgerows, grass strips and reed strips), the area of the three semi-natural habitat types (woodlands, grasslands and wetlands), crop diversity, mean crop field size and all two-way interactions between species identity and the other explanatory variables. All explanatory variables were centered and scaled. Crop type and site identity were added as random effects. We did not include variable ‘year’ in our final models because this variable was never significant and was not relevant to our research question. We accounted for spatial autocorrelation by using an exponential structure on crop field coordinates, and checked for the absence of autocorrelation in the residuals. We used a negative binomial error distribution (type 2: variance increases quadratically with the mean) to deal with over-dispersion. We ran models with a log-link function. We conducted post hoc comparisons of slopes using the emtrends function.

Statistical analyses were run using glmmTMB (Magnusson et al., 2020), entropart (Marcon and Hérault, 2015) and emmeans in R 4.0.5 (R Core Team, 2017).

## 3 Results

The spatial variation in field margin area around the 86 organic crop fields was similar across the three margin types; hedgerows (median = 3.67 ha; range: [0; 17.47]), reed strips (median = 3.60 ha; range: [1.46; 8.72]) and grass strips (median = 3.29 ha; range: [0; 6.27]). The dominant type of semi-natural habitat was grassland (median = 7.38 ha; range: [0; 45.23]), followed by wetland (median = 1.37 ha; range: [0; 48.15]) and by woodland (median = 0.71 ha; range: [0; 20.78]). Crop diversity was on average 0.93 ± 0.04 (median = 0.98; range: [0; 1.6]). Crop mean field size was on average 2.32 ± 0.10 ha (median = 2.27 ha; range: [1.09; 5.85]). There was no correlation among explanatory variables since all Pearson correlation coefficients were under 0.45 (Appendix C).

### 3.1 Forest edge bird guild

Woodland area only had a positive effect on the abundance of one of the nine forest edge species, great tit (*β* = 0.10 ± 0.03, Table 2, Fig. 2), while the area of hedgerows had a positive effect on the abundance of European greenfinch (*β* = 0.15 ± 0.07, Table 2, Fig. 2).

**Table 2.**
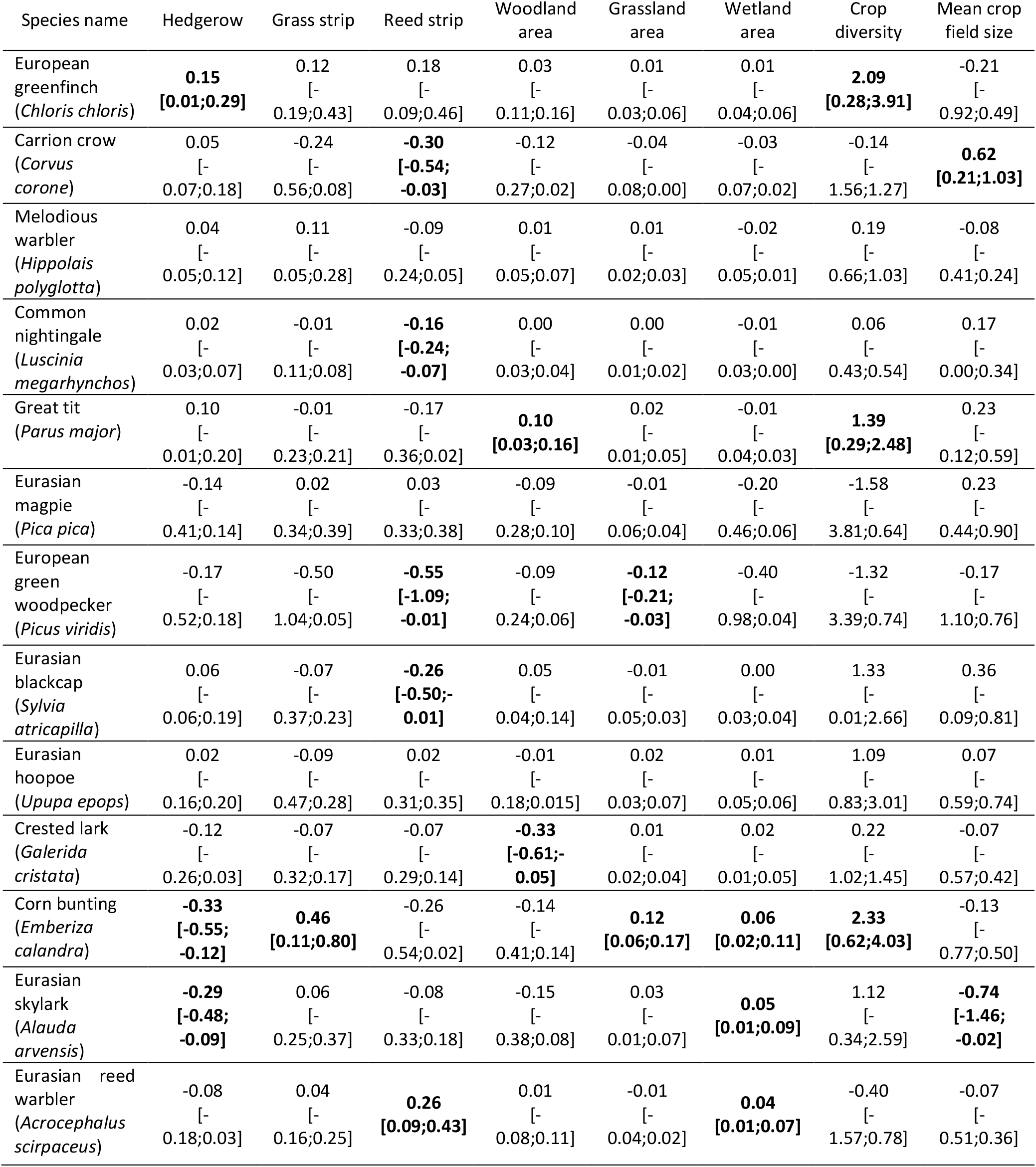
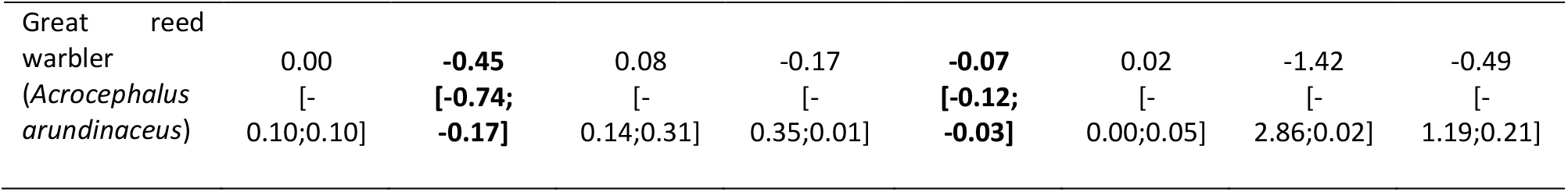
Averaged estimates of the effects of landscape variables for the three bird guilds monitored in agricultural crop fields of the Camargue. The 95 % confidence intervals are in brackets. Values in bold indicate significant effects.

**Figure 2.**
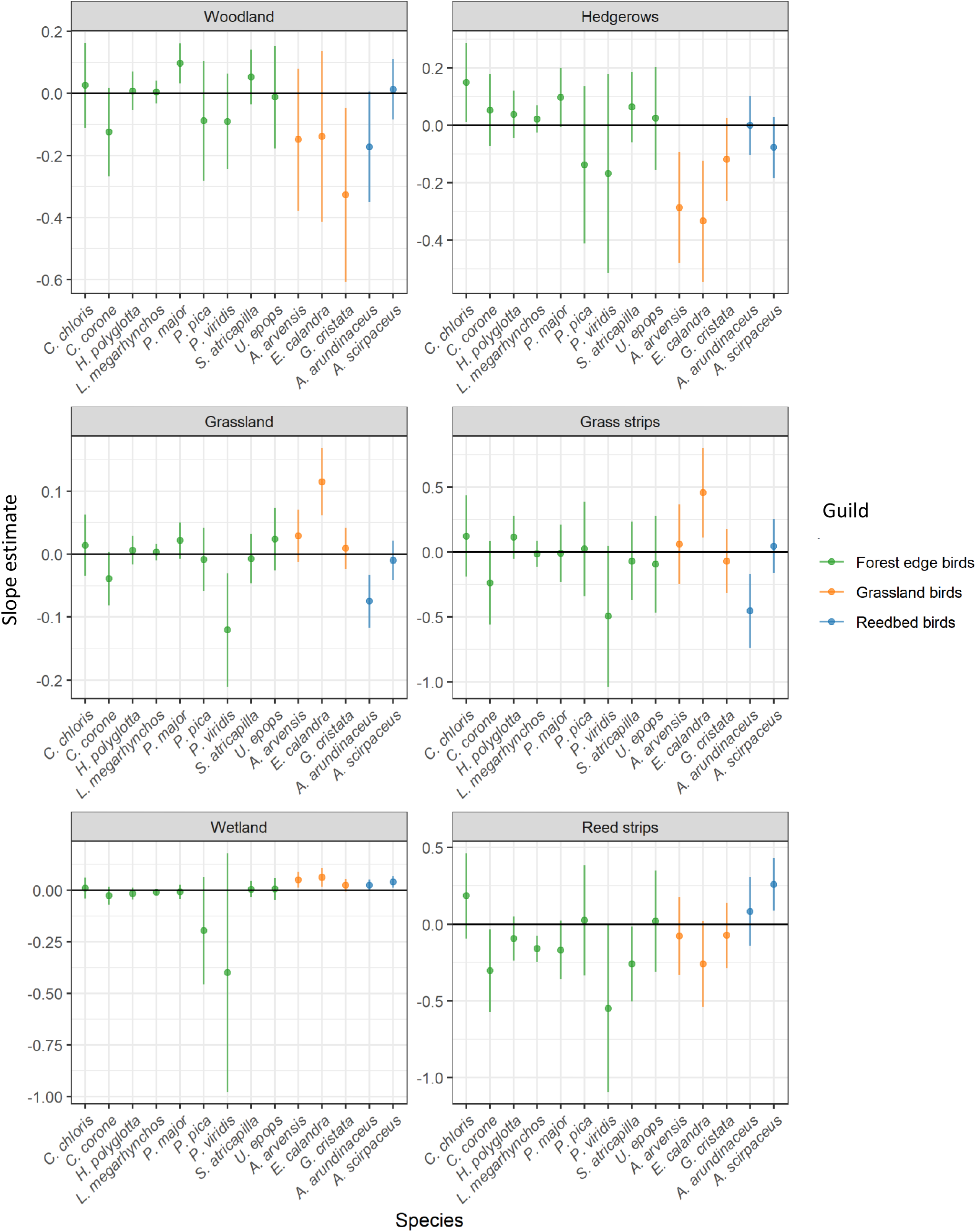
Estimates (± 95% confidence interval) of the effect of landscape variable for each species studied. Each graph corresponds to a landscape variable; the habitat patches on the left and the field margin of the right. The horizontal black line corresponds to 0.. If the 95% confidence intervals does not overlap with zero, the effect of the landscape variable on the abundance of the corresponding species is considered as significant.

Grassland area had a negative effect on the European green woodpecker (*β* = -0.12 ± 0.05, Table 2, Fig. 2).

The area of reed strips had a negative effect on the abundance of carrion crow, common nightingale, European green woodpecker and Eurasian blackcap (respectively *β* = -0.30 ± 0.14, *β* = -0.16 ± 0.04, *β* = -0.55 ± 0.28, *β* = -0.26 ± 0.12, Table 2, Fig. 2).

Crop diversity had a positive effect on the abundance of European greenfinch and great tit (respectively *β* = 2.09 ± 0.92, *β* = 1.39 ± 0.56, Table 2), while crop mean field size had a positive effect on the abundance of carrion crow (*β* = 0.62 ± 0.20, Table 2).

There was no significant effect of wetland area or grass strip area on the abundance of species of this guild (Table 2, Fig. 2).

### 3.2 Grassland bird guild

The abundance of corn bunting was positively related to both grassland area (*β* = 0.12 ± 0.03, Table 2, Fig. 2) and the area of grass strips (*β* = 0.46 ± 0.18, Table 2, Fig. 2).

Woodland area had a negative effect on the abundance of crested lark (*β* = -0.33 ± 0.15, Table 2, Fig. 2), while the area of hedgerows had a negative effect on the abundance of Eurasian skylark and corn bunting (respectively *β* = -0.29 ± 0.10, *β* = -0.33 ± 0.11, Table 2, Fig. 2).

Wetland area had a positive effect on the abundance of Eurasian skylark and corn bunting (respectively *β* = 0.05 ± 0.02, *β* = 0.06 ± 0.02, Table 2, Fig. 2), while the area of reed margins had no effect on the abundance of grassland species (Table 2, Fig. 2).

Crop diversity had a positive effect on the abundance of corn bunting (*β* = 2.33 ± 0.87, Table 2), while crop mean field size had a negative effect on the abundance of Eurasian skylark (*β* = -0.74 ± 0.37, Table 2).

### 3.3 Reedbed bird guild

The abundance of Eurasian reed warbler was positively related to both wetland area (*β* = 0.04 ± 0.01, Table 2, Fig. 2) and the area of reed margins (*β* = 0.26 ± 0.09, Table 2, Fig. 2).

The abundance of great reed warbler was negatively related to both grassland area (*β* = - 0.07 ± 0.02, Table 2, Fig. 2) and the area of grass strips (*β* = -0.45 ± 0.14, Table 2, Fig. 2).

There was no significant effect of woodland area, hedgerow area, crop diversity or crop mean field size on the abundance of species of this guild (Table 2, Fig. 2).

## 4 Discussion

Our study shows that different types of field margins can provide alternative habitats to terrestrial birds in a rice paddy landscape, but species responses vary even within species guilds. We found that (i) grass strips represent a substitute habitat to grasslands for corn bunting and (ii) reed strips represent a substitute habitat to wetlands for the Eurasian reed warbler, in line with the habitat compensation hypothesis. For these two species, the positive effect of field margins on the abundance of species was even stronger than the effect of the corresponding semi-natural habitat patch. This result suggests that field margins are currently valuable habitat rather than substitute ones for these two species. It is consistent with the meta-analysis conducted by Riva and Fahrig (2022), which highlighted the higher value of small habitat patches for biodiversity conservation. In contrast, we could not confirm the compensation hypothesis for 12 out of 14 species. Such a lack of support to the compensation hypothesis could be explained by different methodological and ecological reasons. First, we observed a general lack of species responses to their primary habitat with only 3 species responding positively to the primary habitat surface area. This may result from the use of broad categories of habitat preferences, while species abundance may vary along ecological continuums. Also, semi-natural habitats have been grouped into three primary habitat categories, which may not be detailed enough to match species habitats preferences. For example, wetlands include reedbeds but also ponds without emergent vegetation which are likely not very attractive for reedbeds birds. A more detailed mapping of primary habitats or functional description of habitats, such as habitat quality, nesting opportunities or food resources would therefore be necessary to further test the habitat compensation hypothesis for several of the species considered. In addition, the observed species might potentially accommodate a diversity of habitats. Indeed, in the Camargue, some forest edge species like carrion crow, Eurasian magpie or common nightingale are known to be able to nest in very open landscape e.g. in isolated trees within a matrix of cultivated fields. Further studies aiming to test the habitat compensation hypothesis should therefore focus on species that are more strongly associated with their primary habitat.

Our results show that the compensation hypothesis cannot be generalized to all bird species within the three guilds studied. Indeed, only some species benefited from the presence of field margins as substitute habitat. Moreover, some species within these guilds were not even recorded within sampled agricultural landscapes. For example, the bearded reedling (*Panurus biarmicus*), a reedbed bird, the blue tit (*Cyanistes caeruleus*), a forest edge bird, or the tawny pipit (*Anthus campestris*), a grassland bird, breed in Camargue but were not contacted at all during our surveys.

The lack of effect of field margins on some species may be partly explained by both the quality of field margins and the ecological preferences of these species. Indeed, in Camargue, ditches are increasingly being lined with concrete or buried, like in Japanese rice paddy landscapes for example (Yamada et al., 2011). Some studies have highlighted that earthen ditches host much more aquatic fauna and flora than concrete ones (Katoh et al., 2009). It was also shown that the density of intermediate egrets (*Egretta intermedia*) was twice as high in rice fields with shallow earthen ditches than in rice fields with deep concrete-lined ditches (Katayama et al., 2012). Here, we found a positive effect of reed field margins for the Eurasian reed warbler but not for the great reed warbler, the latter requiring wetter and larger patches of reedbeds than the Eurasian reed warbler (BirdLife International, 2022). The absence of the bearded reedling is also consistent with the fact that this species requires larger areas of reedbeds and is not encountered in reed strips along artificial ditches (P.M. pers. obs.).

Our results nearly support the hypothesis that hedgerows represent a substitute habitat for great tit with a positive effect of woodland and a positive but no significant effect of hedgerow. The European greenfinch is the only species significantly positively affected by hedgerow, a result that may be useful to encourage farmers to maintain and restore hedgerows. Yet, the lack of effect of hedgerows for the other species was surprising since hedgerows are known to benefit a broad range of forest edge species (Batáry et al., 2010; Wilson et al., 2017). In Camargue, the poor quality of hedgerows may explain the lack of response within a wider bird community because several of them, i.e. coniferous or giant cane (*Arundo donax*) hedgerows, are not suitable to forest edge birds as their volume and plant diversity are low (Graham et al., 2018; Montgomery et al., 2020).

Our results highlight that grass strips have a stronger effect than grasslands for corn bunting. The greater plant biomass of grass strips compared to Mediterranean salt meadows, which constitute most of the grassland area habitat category, may explain this greater effect of grass strips compared to other open habitats. The high density of seeds available in cultivated fields where this species comes to feed (Madge and de Juana, 2020), can also be a confounding effect. Unlike other types of field margins, grass strips are probably not used as a nesting habitat due to disturbance from agricultural activity. In particular, these strips are frequently mowed and used by farmers to move around the crop fields, which causes disturbances that might prevent nesting (Vickery et al., 2009).

Further research should therefore assess the ecological value of field margins, for instance by comparing the demographics of Eurasian reed warbler and corn bunting in the substitute habitat and in natural habitat to ensure that field margins are not ecological traps (Horne, 1983). This would also allow to develop recommendations on the most favorable field margin management methods. It may also be relevant to study the role of different types of field margins for generalist species. Indeed, a recent paper has highlighted the progressive colonization of farmland habitats by generalist bird species over the last decades in Spain (Díaz et al., 2022). Taking into account the response of generalist bird species may therefore help avoiding the homogenization of bird communities in rice paddy landscapes. Finally, the value of field margins may also depend on the availability of habitat patches within the landscape. For instance, reedbeds may have a more positive effects when they are close to a large patch of wetland. Testing such interactive effects would require an adequate study design with all combinations of values for field margins and semi-natural patches, and a sample size large enough to provide robust estimates of all parameters within associated statistical models.

Our study also highlights that increasing a type of field margins may have antagonistic effects across different guilds. Indeed, four species within the forest edge bird guild were negatively impacted by the area of reed strips. This result may be due to the fact that this type of field margin provides too few resources in terms of food and nesting sites for forest edge bird species (Shoffner et al., 2018). Similarly, the abundance of the great reed warbler is negatively correlated to the area of grassland and grass strip as this species occur mainly in wet habitats during the breeding season (Dyrcz, 2020). As expected, grassland birds were negatively impacted by the area of hedgerows and woodland confirming previous studies that observed a similar negative effect of wooded habitats on different species of grassland birds (e.g. Ellison et al., 2013; Wilson et al., 2014). Woodland patches usually do not offer resources for grassland birds and are avoided because they are a source of avian and mammalian predators (Burger et al., 1994). Our study therefore confirms that it may not be possible to favor all bird species within a single landscape and it may be necessary to focus on the type of field margins that most favor species in need of conservation attention.

Our study confirms that increasing crop diversity and decreasing crop mean field size are complementary levers to promote biodiversity in agricultural landscapes (Sirami et al., 2019). Indeed, crop diversity benefited two of the nine forest edge species, European greenfinch and great tit and one grassland species, corn bunting. Moreover, the decrease in crop field size had a positive impact on Eurasian skylark. The results likely stem from the fact that higher landscape heterogeneity provides readily available complementary resources (Batáry et al., 2017; Sirami et al., 2019). On the other hand, we found a positive effect of the increase in crop field size on the abundance of carrion crows. This effect is probably related to the fact that this species feed in groups on the ground and may thus be favored by large open areas (Madge, 2020).

In conclusion, our results highlight that field margins are valuable landscape components to improve biodiversity conservation but cannot be the only components to be promoted in rice paddy landscapes. In Camargue, current conservation priorities concern the disappearance of wetlands and grasslands as well as the degraded conservation status of species associated with these habitats, whereas there is less concern for forest edge birds, which can be found in other agricultural landscapes. Our study therefore suggests that conserving and restoring wetlands and grasslands and the associates field margins, reed strips and grass strips, represent a promising avenue to increase biodiversity in the agricultural landscapes of Camargue. On the other hand, despite the negative impact of hedgerows on grassland birds and waterbirds (Tourenq et al., 2001), they can host a diversity of auxiliary species as well as taxa of high conservation importance in Camargue and other wetlands such as bats (Mas et al., 2021). Hedgerows have also been shown to limit the presence of greater flamingos (*Phoenicopterus roseus*), considered as a pest in rice fields (Ernoul et al., 2014). Taking into account the role of hedgerows across taxa would be particularly relevant in the context of the current action plan of replanting hedgerows carried out locally by the Regional Natural Park of the Camargue. Land-use planning studies could be a good way to propose management actions to farmers and stakeholders, maximizing both long-term agricultural benefits and biodiversity conservation.

## Acknowledgements

We are grateful to all farmers and landowners who graciously permitted us to work in their fields. We are particularly grateful to the company Biosud who helped with setting up the network of organic farmers where we sampled avian biodiversity. Finally, particular thanks to Fabien Laroche for his advices on statistical analysis.

## Data, script and code availability

Supplementary information, dataset and statistical scripts are available here: https://doi.org/10.5281/zenodo.7685771

## Supplementary information

Appendix A: Map of habitat localization in Camargue

Appendix B: Table of species guilds

Appendix C: Correlation table between landscape explanatory variables and sampling completeness (Cn).

