## Supplementary material for "Field margins as substitute habitat for the conservation of birds in agricultural wetlands"

### Appendix A: Habitat localization in Camargue

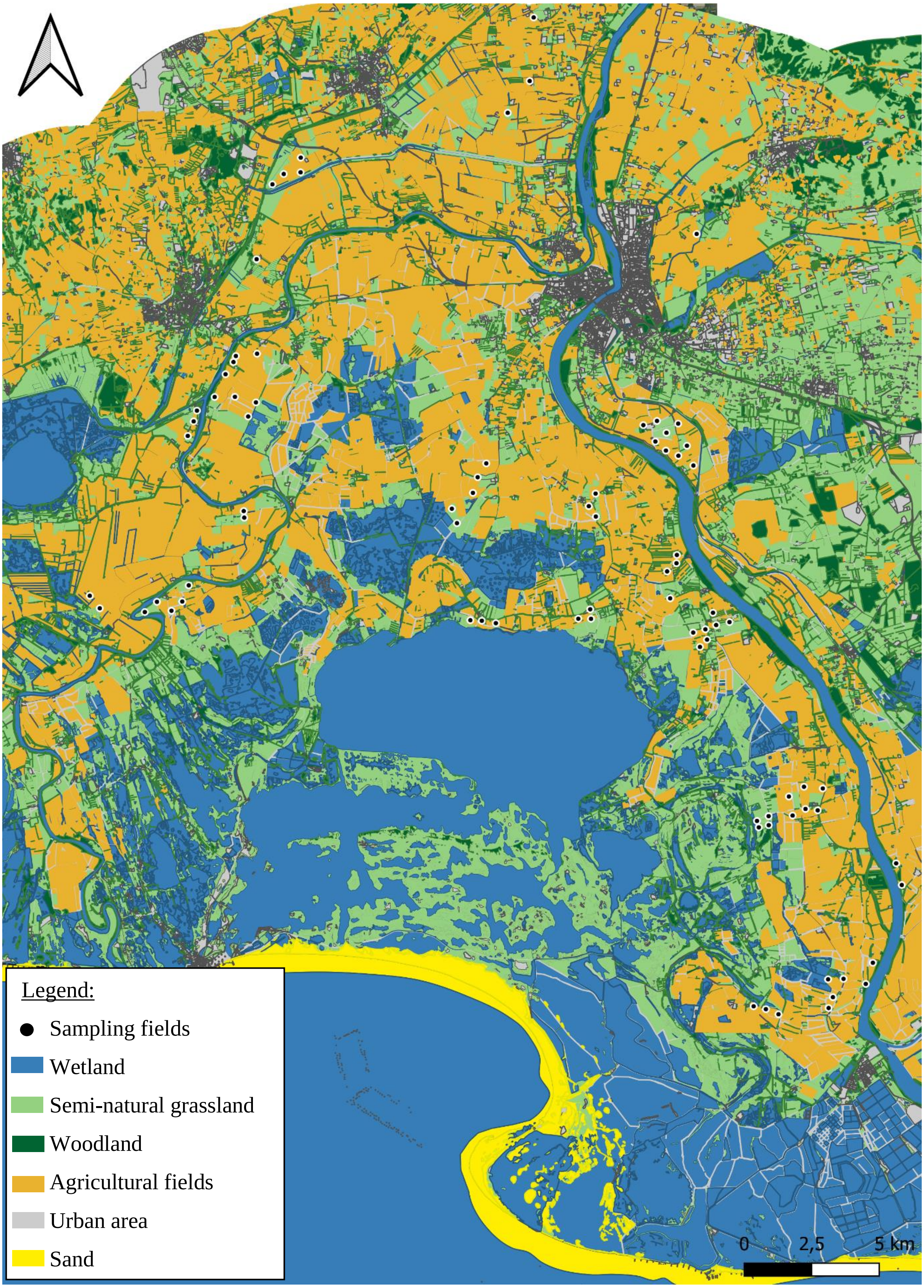

Appendix B: Eunis reproductive habitat classification and IUCN status for the species present in more than 15 % of the sampled crop fields. Categories B, C and D (costal habitats, inland surface waters and mires, bogs and fens) are related to wetland habitats used by reedbed birds. Categories E and I (grasslands and lands dominated by forbs, mosses or lichens and regularly or recently cultivated agricultural, horticultural and domestic habitats) are related to grassland habitats used by grassland birds. Categories F and G (heathland, scrub and tundra and woodland, forest and other wooded land) are related to wooded habitats used by forest edge birds. Category J (constructed, industrial and other artificial habitats) are related to urban habitat. Urban species and species in more than one habitat category were not assigned to one of our three guilds. See the Eunis habitat classification (Davies et al., 2004) for details of the different habitat types and sub-categories X (habitat complexes). Common pheasant (*Phasianus colchicus*) was not included in the analysis due to the large number of breeding birds released all year round for hunting. Their presence could therefore be biased by the release site.

| Species | Eunis habitat code | Experts opinion | Bird guild |
|---|---|---|---|
| *Acrocephalus arundinaceus* | C2-3, X01-03 | Agreed | Reedbed |
| *Acrocephalus scirpaceus* | C1-3, D5 | Agreed | Reedbed |
| *Alauda arvensis* | E2-3, I1.1 | Agreed | Grassland |
| *Cettia cetti* | F9, G1.1-1.3 | Species locally link to woodland and wetland | Generalist |
| *Chloris chloris* | G1-2-3-4-5, FA, X10-11-23-24-25 | Agreed | Forest edge |
| *Cisticola juncidis* | C3, E1-2-3 | Agreed | Generalist |
| *Columba palumbus* | FA, G, X10-11 | Agreed | Generalist |
| *Corvus corone* | G1-2-3-4-5, X10-11 | Agreed | Forest edge |
| *Cuculus canorus* | F3, G1-2-3-4-5, X | Could parasite *accrocephalus ssp.* nest, thus species also link to wetland | Generalist |
| *Emberiza calandra* | B1, E2-3-6-7, F4, I1 | Only linked to grassland in the Camargue | Grassland |
| *Galerida cristata* | B1, C2, F6, FB.4, I1.3-1.5, J1 | Only linked to grassland in the Camargue | Grassland |

| | | | |
|---|---|---|---|
| *Hippolais polyglotta* | FA, G5.6-5.7-5.8-1.C, I1.5, J3.3-6 | No artificial habitat in the study area thus strictly link to woodland | Forest edge |
| *Luscinia megarhynchos* | F5-6, FA, G1-2-5, I2, X11 | Agreed | Forest edge |
| *Motacilla flava* | C3, E2-3, X02-03 | Agreed | Generalist |
| *Parus major* | FA, G1-2-3-4-5, X10-11-22-23-24 | Agreed | Forest edge |
| *Passer domesticus* | J1-2 | Agreed | Urban |
| *Phasianus colchicus* | - | - | - |
| *Pica pica* | FA, FB, G, X10-11 | Agreed | Forest edge |
| *Picus viridis* | FA, G1-2.1-5, X10 | Agreed | Forest edge |
| *Streptopelia decaocto* | F5-6, FA, G1-5, J1-2, X11 | Agreed | Generalist |
| *Sturnus vulgaris* | G1-2-4, J1, X10-11 | Agreed | Generalist |
| *Sylvia atricapilla* | F, G, I, X06-10-11-20-22-23-24-25 | Only linked to woodland in the Camargue | Forest edge |
| *Upupa epops* | FB.4, G1.D-2-2.9-3-5.8, J2, X10-11 | Agreed | Forest edge |

18

Appendix C: Correlation table between landscape explanatory variables and sampling completeness (Cn).

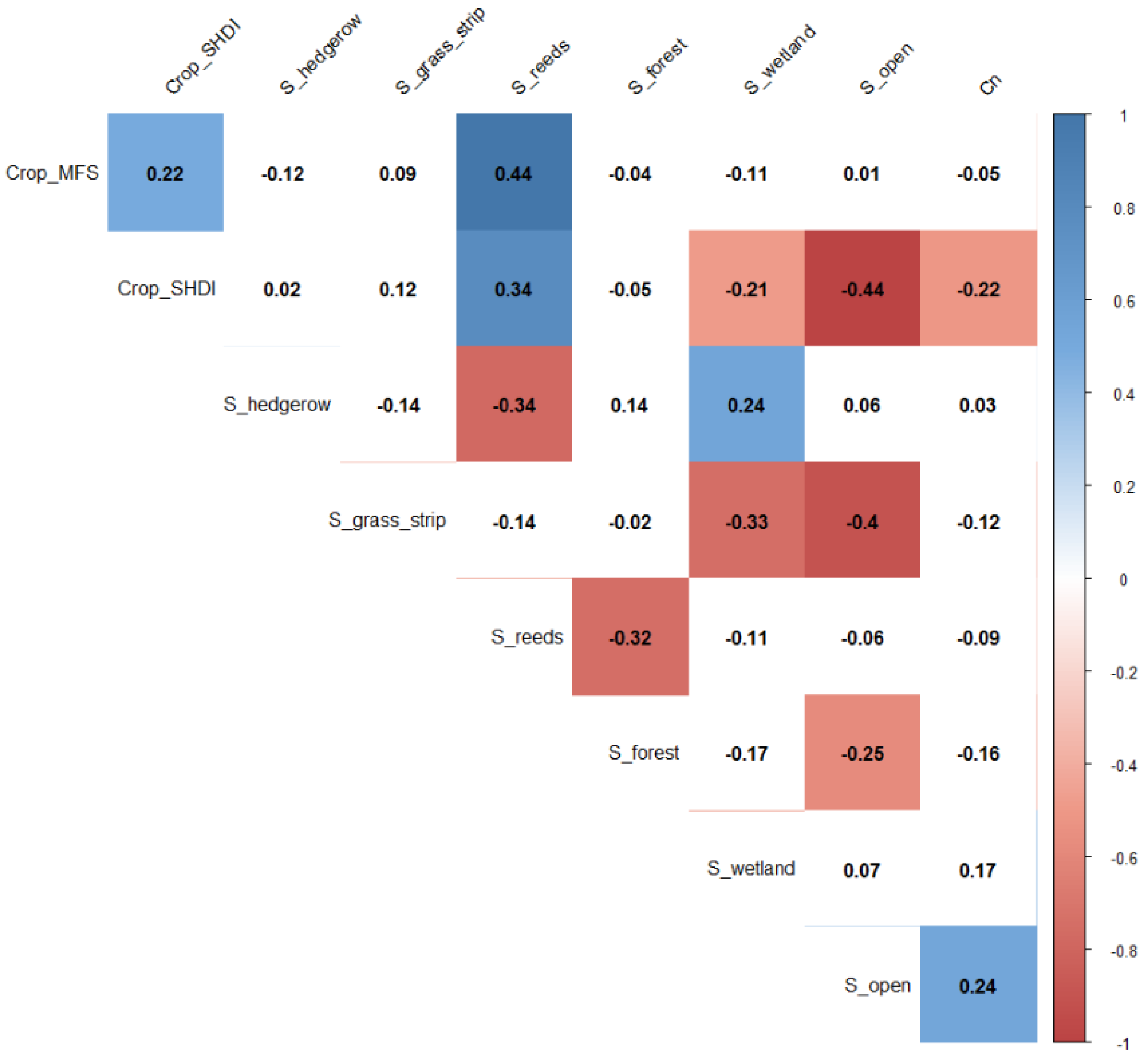
